# Diurnal activity budget and conservation implication of *Hippopotamus amphibius* at Bui National Park, Ghana

**DOI:** 10.1101/2021.08.30.458240

**Authors:** Godfred Bempah, Marie Osei, Alice Gyimah, Winnifred Nsoah

## Abstract

Animals apportion time for their daily behavioral activities. We studied the activity budget of *Hippopotamus amphibius* at the Black Volta River in the Bui National Park, Ghana. We performed instantaneous scan sample using ground count survey between August 2020 to July 2021. We observed that *H. amphibius* spent most of their day time resting (54.75 %), followed by feeding (22.93 %), walking (19.2 %) and touching (3.12 %). We found significant difference in the time spent between the daily activities (H = 41.67; *p* < 0.0001). Except activities involving touching, we found no significant seasonal differences in feeding, resting and walking activities by *H. amphibius*. Understanding the behavioral activities of *H. amphibius* will enhance management and conservation of the animals.

## 1. Introduction

Animals respond differently to stimuli in their environment and this is referred to as behavior [1]. Animals’ ability to find food and water, as well as noticing a mate, predator or competitor constitutes such stimuli in the environment. Any alteration to an animal’s habitat is likely to have an effect on the individuals and subsequently the entire population. To meet the daily requirements for survival and reproduction, animals apportion time for their daily behavioral activities [2]. One of such species that exhibit different behaviors attributed to changes in their environment is the *Hippopotamus amphibius* (hippopotamus).

The hippopotamus, which is a semiaquatic species is among the mega fauna species of the world. It utilizes two basic habitats including resting in water at day time and grazing on sward during night time [3]. It plays important ecological role in water and terrestrial ecosystems by enhancing biological diversity of their environment [4-5]. At sexual maturity, male and female hippopotamus mate in water and after 8 months of gestation, the female gives birth mostly in water with lower depth [6]. The hippopotamus aggregate in groups due to irregular distribution caused by the patchy distribution of their feeding and water resources, allowing them to improve their reproduction success [7].

The feeding behavior of hippopotamus enable them to consume an amount of 35-50 kg of grass daily [8]. The hippopotamus feed relatively in a solitary manner but when in water they come together and create a bond. This behavior however could change when there is any change in the resources they rely on. In the Bui National Park (BNP), which harbors a sizable population of hippopotamus in Ghana, the animals spend most of their time and activities in water resources [9]. As a results of several factors including anthropogenic effects, water and feeding resources have changed in the BNP causing the hippopotamus to react and adjust their behavior to the varying conditions [2].

According to Bateson and Martin [10], activity budget refers to the amount of time an animal apportions to a particular behavioral activity. The diurnal behavioral patterns of the hippopotamus involve behavioral activities and behavioral events. Behavioral activities have a longer period of occurrence including resting, feeding, walking etc. while behavioral events are shorter including barking, yawning, threats etc [2]. To effectively conserve and manage hippopotamus, it is important to understand the changes in the animals’ behavior, especially during changing conditions in the environment. The study did not give detail consideration to the mating behavior of the animals due to the unpredictable aggressiveness of the hippopotamus which could cause harm to members of the research team. Furthermore, we only considered behavioral activities but not behavioral events of the hippopotamus due to lack of resources.

There exists a gap in literature on changes in hippopotamus’ behavioral activities caused by human mediated land-use especially relating to the animal’s presence in the Black Volta River (BVR) of the BNP where a hydroelectric dam has been constructed recently. Therefore, this study aims to establish the behavior of hippopotamus during the wet and dry seasons. We hypothesized that; 1) The hippopotamus spends more time resting than other activities to protect its skin from sun heat, and 2) There is seasonal differences in the amount of time spent for the various behavioral activities by the hippopotamus.

## 2. Materials and Methods

### 2.1. Study area

The study was carried out along the BVR in BNP (80 23’ 13.2072” N, 20 22’ 43.9788” W). The BNP is the third largest protected area in Ghana with a size of about 1821 km2 (Fig. 1). It is one of the few remaining habitats of hippopotamus in Ghana and has very rich flora and fauna resources. The BNP is located within the Bono and Savanna regions of Ghana.

**Figure 1:**
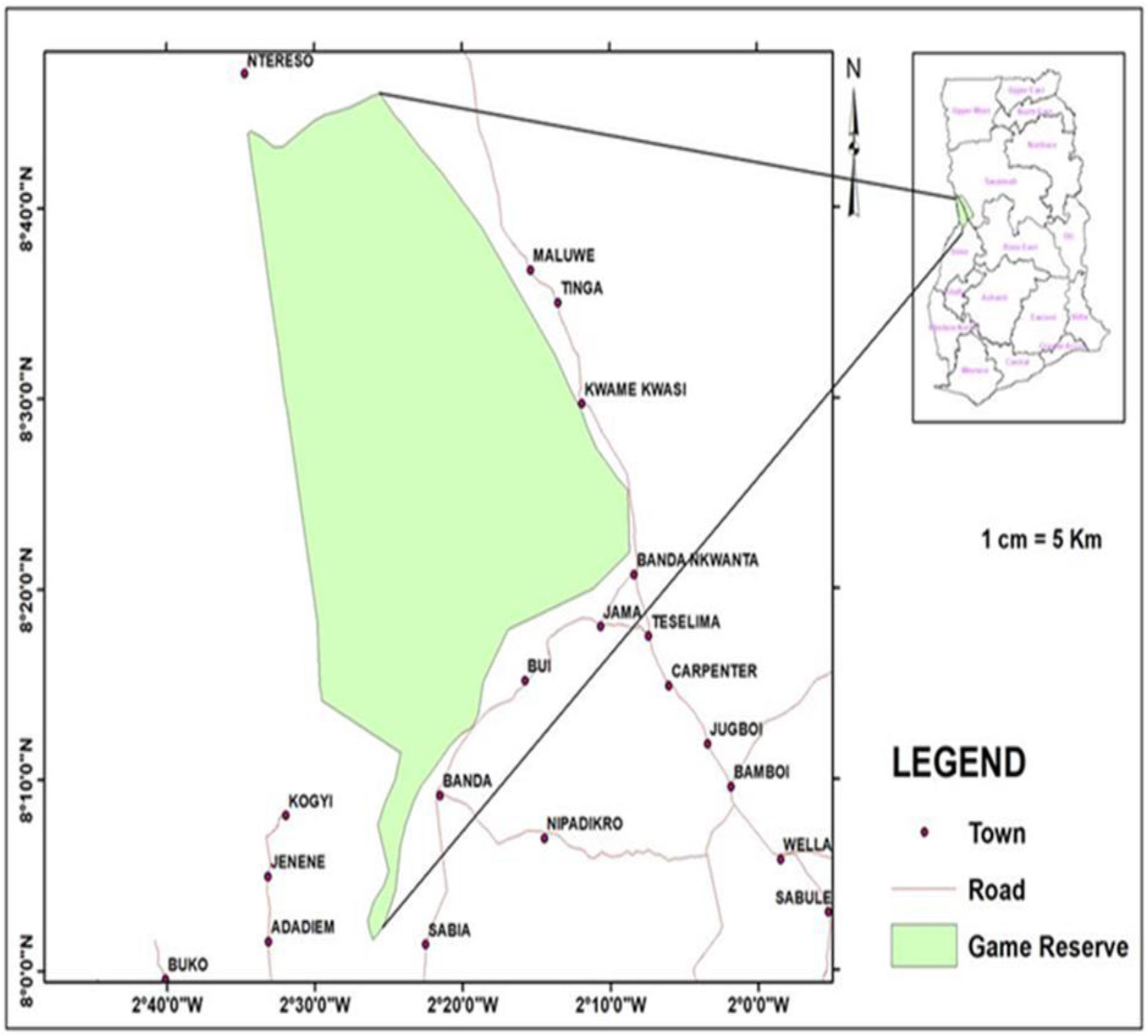
Map of Bui National Park, Ghana

### 2.2. Data recording

We observed and recorded hippopotamus activities twice each month with two weeks’ time interval for a period of 12 months from August 2020 - July 2021, daily observation time between 06: 00 am - 06: 00 pm during the dry season (December – March) and wet season (April – November). We followed the hippopotamus using a canoe at a distance of 30 m to avoid disturbing the animals as well as ensuring that we are secured from animal attacks. Animals were observed directly and with the aid of binoculars. When water was so low to use a canoe, we walked at the river-shore.

We used instantaneous scan to observe the animals. Daily study period was divided into four blocks: 6:00 am - 9:00 am, 9:00 am - 12:00 pm, 12:00 pm - 3:00 pm, 3:00 pm - 6:00 pm. Within each block period, we spent 30 mins to observe animals for 5 mins at every 7 mins time interval. In total, we spent three hours (hrs) during daily scans at the site per month totaling 216 hours (3 hrs * 2 days * 12 months * 3 sites = 216 hrs). We recorded by scanning behavioral activities applying methods by Bateson and Martin [10] and recorded the time spent on activities including resting, walking, feeding and touching.

### 2.3. Data analysis

We summarized data using Microsoft excel and determined any statistically significant seasonal difference between the behavioral activities using Mann-Whitney U and Kruskal Wallis tests. Statistical analysis was performed using Paleontological Statistics (PAST) software package PAST version 3.0 [11].

## 3. Results

### 3.1. Diurnal activity budget of hippopotamus

The hippopotamuses were observed to spend most of their day time resting (54.75 ± 1.22%), followed by feeding (22.93 ± 0.64%), walking (19.2 ± 0.92 %) and touching (3.12 ± 0.4%) constituting their daily behavioral activities (Fig. 2). We found a significant difference in the time spent between the daily activities (H = 41.67; *p* < 0.0001).

**Figure 2:**
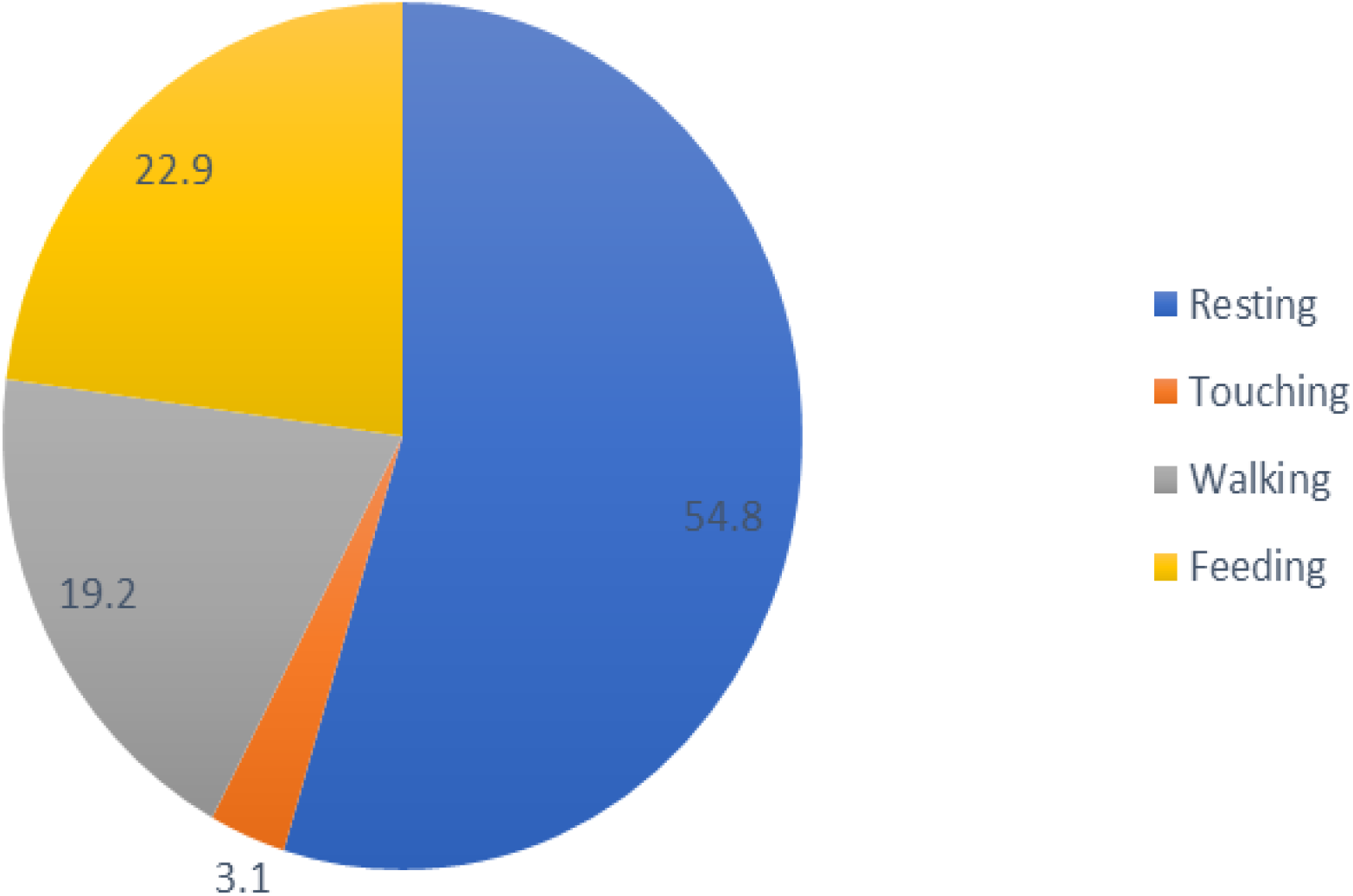
Activity budget of the hippopotamus during the study period at Bui National Park

### 3.2. Seasonal differences in activity budget

We found that hippopotamus spend significant amount of time on touching in the dry season than wet season (U = 4; *p* = 0.047) in the Bui National Park (BNP). Feeding activities were higher in the dry season (24.15 %) than wet season (22.32%). Resting activities were higher in the wet season (56.19%) than dry season (51.87%, Fig. 3). We found a no significant difference between the wet and dry seasons for activities of resting (U = 7.5; *p* = 0.165), walking (U = 14; *p* = 0.808) and feeding (U = 8; *p* = 0.214).

**Figure 3:**
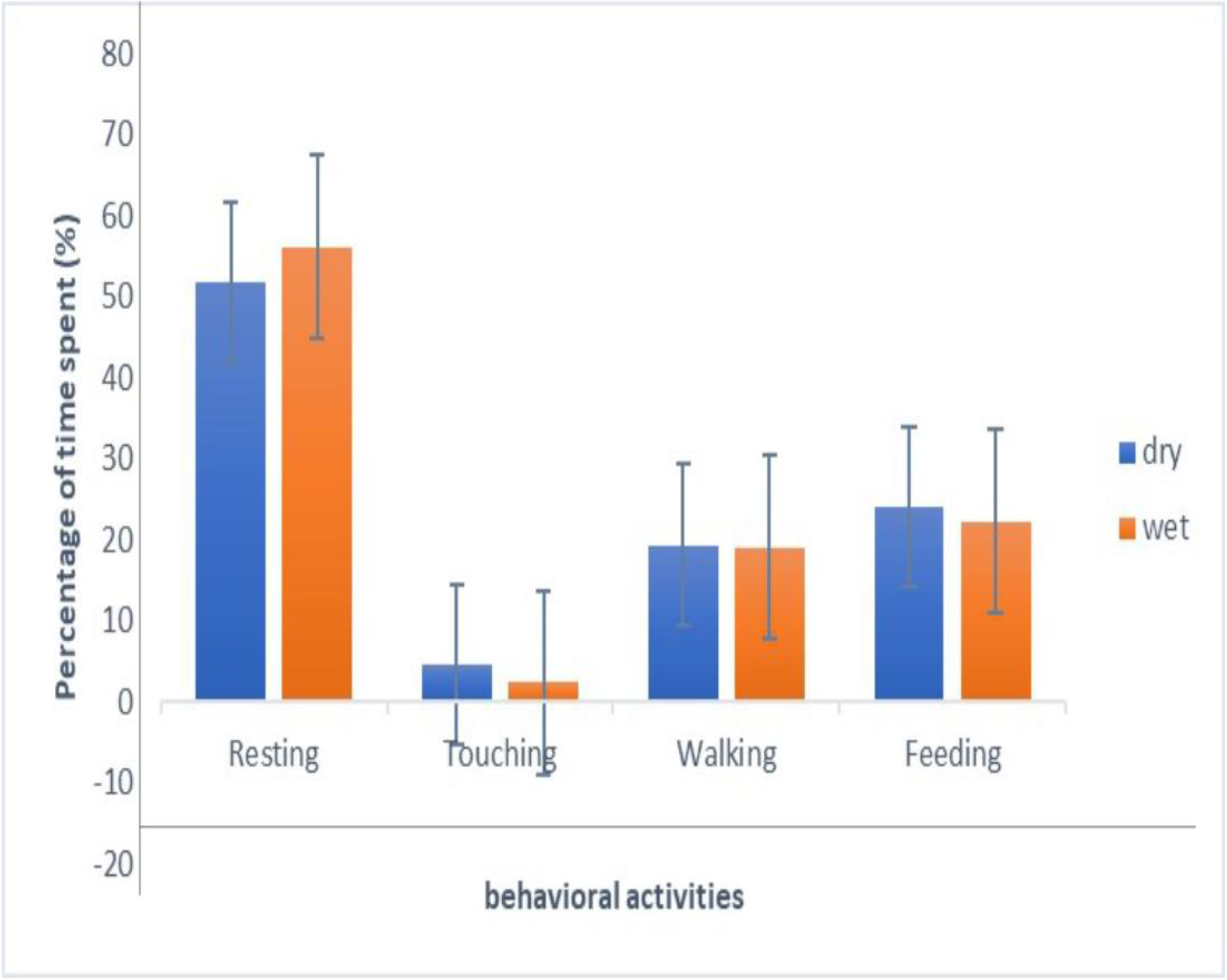
Seasonal differences in the percentage of time spent on the various behavioral activities by the hippopotamus at Bui National Park

### 3.3. Differences in monthly activity budget

Resting was highest in June with 60.9 ± 1.3% and lowest in March (47.2 ± 0.8%), while feeding activities were highest in March (25.9 ± 0.6%) and lowest in December (19.7 ± 0.5%), touching was highest in December (6.1 ± 0.2%) and lowest in August (1.1 ± 0.1%), and walking activities were observed to be highest in September (22.9 ± 0.1%) and lowest in January (12.3 ± 0.3%, Table 1).

**Table 1:** Mean monthly recordings of the activity budget of hippopotamus from August 2020 - July 2021 at Bui National Park

| Month | Resting | Touching | Walking | Feeding |
| --- | --- | --- | --- | --- |
| August | 58.3 ± 0.8 | 1.1 ± 0.1 | 18.7 ± 1.2 | 21.9 ± 0.2 |
| September | 51.4 ± 1.5 | 1.9 ± 0.2 | 22.9 ± 0.1 | 23.8 ± 0.4 |
| October | 58.8 ± 0.2 | 1.2 ± 0.1 | 17.6 ± 0.5 | 22.4 ± 0.6 |
| November | 55.7 ± 0.4 | 2.3 ± 0.2 | 18.3 ± 0.4 | 23.7 ± 0.2 |
| December | 52.4 ± 0.6 | 6.1 ± 0.2 | 21.8 ± 0.6 | 19.7 ± 0.5 |
| January | 56.5 ± 1.5 | 4.1 ± 0.1 | 12.3 ± 0.3 | 27.1 ± 0.2 |
| February | 51.4 ± 1.4 | 3.5 ± 0.5 | 21.3 ± 0.7 | 23.9 ± 0.2 |
| March | 47.2 ± 0.8 | 4.6 ± 0.4 | 22.3 ± 0.4 | 25.9 ± 0.6 |
| April | 55.2 ± 1.1 | 5.1 ± 0.3 | 19.9 ± 0.1 | 19.8 ± 0.4 |
| May | 50.1 ± 0.9 | 3.85 ± 0.35 | 22.4 ± 0.5 | 23.6 ± 0.9 |
| June | 60.9 ± 1.3 | 2.1 ± 0.2 | 15.7 ± 0.3 | 21.3 ± 0.4 |
| July | 59.1 ± 1.8 | 1.5 ± 0.1 | 16.95 ± 0.95 | 22.1 ± 0.2 |

### 3.4. Allocated daily time for activities

Figure 3 shows the percentages of time spent by hippopotamus for various activities at different time of the day. Feeding and walking activities peaked in the early morning (06:00 am - 9:00 am) and in the late afternoon (3:00 pm - 6:00 pm). Resting only peaked in late morning (06:00 am - 9:00 am) with 44% and showed constant trend for the rest of the day. Touching was also high in the late morning (06:00 am - 9:00 am) with 25.5% (Fig 4).

**Figure 4:**
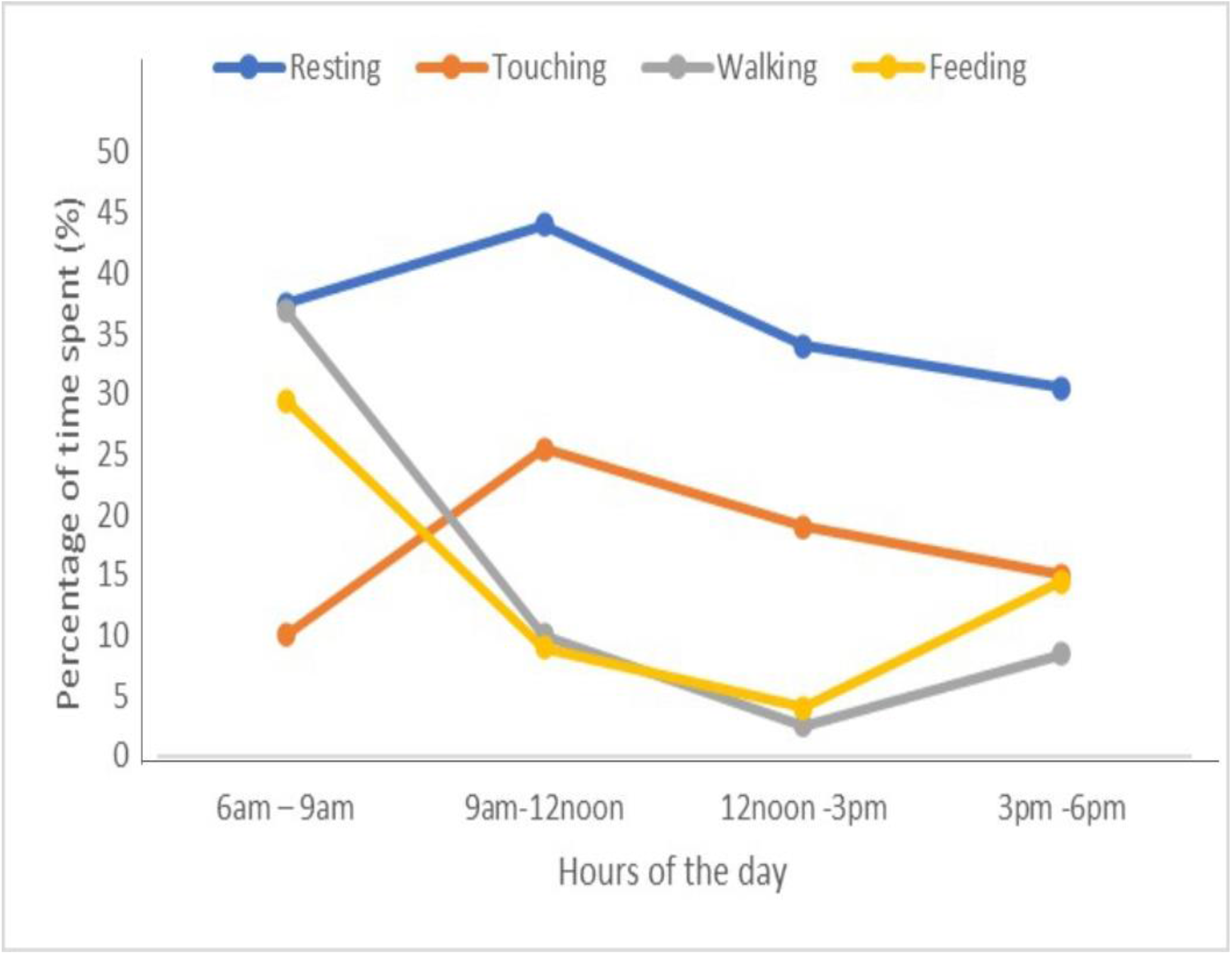
Percentages of time spent by hippopotamus to perform behavioral activities at different time of the day at Bui National Park

## 4. Discussion

Animals apportion time for their daily behavioral activities. We observed, hippopotamus spent about 54.8 % of their day time resting while almost half of the remaining day time was spent feeding, walking and touching. More feeding activities were performed in the dry season whiles resting was more prevalent in the wet season. However, we also found no seasonal difference in resting, feeding, walking, while touching activities among hippopotamus was seasonally different.

The allocation of time by hippopotamus for their daily activities is similar to results in other studies on hippopotamus [2,12]. This confirms that hippopotamuses are more active at night than day time [6]. We observed that hippopotamus spent almost the same amount of time for both walking and feeding, even though feeding was slightly higher as corroborated in other studies [2]. Feeding and walking had similar trend throughout the day. Much feeding and walking activities were performed by hippopotamus in the early morning and gradually with the progression of the day, they reduce feeding and walking, and rest under tree shade or move into the water while touching each other.

During sunny hours of the day, air temperature increases and this propels hippopotamus to rest in water or under tree shades. hippopotamus adopt this strategy to keep their skin from direct sun heat through thermoregulation [13]. The hippopotamus feed in the remaining hours of the day only during the wet season or cloudy atmosphere [12]. Seasonality affected touching among the hippopotamus as they touched themselves more in the dry season perhaps to maintain the social bond when the amount of water was limited and increased air temperature. We observed that during afternoons, hippopotamuses were mostly submerged in the water, similar to results by Saikawa et al. [14], but sometimes came out of the water to stand or rest under tree shades and moved back into the water [2].

Seasonality did not strongly affect the activity budget of the hippopotamus as we observed no significant differences between wet and dry seasons for feeding, walking and resting activities. This means that hippopotamus maintained similar amount of time spent on the various behavioral activities for both wet and dry seasons. Feeding and walking assumed almost the same amount of time. The hippopotamus mostly walked to search for food. Therefore, the proximity of available food resources to the water most likely will reduce the amount of time spent on walking by hippopotamus [15]. We found a significant difference in the amount of time spent touching by hippopotamus between wet and dry seasons with more touching observed in the dry season than wet season. Our observation is in agreement with the findings of Timbuka [2]. This could be attributed to hippopotamus forming bonds during periods of low water availability.

The hippopotamus is noted to graze at night usually spending 12 hrs on feeding [6], however our observation that hippopotamus grazed in the early mornings of the day could possibly be as a results of low food resources availability in the Bui National Park (BNP), thus the animals acquired less forage at night, hence more grazing to satisfy nutritional needs and maximize intake of energy [16]. It could also be that they favored the grazing resources available at that specific area and particular time [2]. The quality and quantity of resources used by hippopotamus most likely affect the diurnal activity budget of hippopotamus [17] and hence the amount of time spent on the various behavioral activities.

## 5. Conclusion

We observed that hippopotamus spent most of their daily time resting. Feeding mostly occurred in the early morning and was simultaneously performed with walking activities. We did not observe seasonal differences in the amount of time spent performing the various behavioral activities except for touching. This study is important for wildlife managers to understand the various daily behavioral activities of the hippopotamus and adopt measures for the animals’ conservation.

## Acknowledgement

We wish to thank the staff of Bui National Park and the local people residing along the Black Volta River for their support during the field work.

